# Intraspecific size variation in planktonic foraminifera cannot be consistently predicted by the environment

**DOI:** 10.1101/468165

**Authors:** Marina C. Rillo, C. Giles Miller, Michal Kucera, Thomas H. G. Ezard

## Abstract

The size structure of plankton communities is an important determinant of their functions in marine ecosystems. However, few studies have quantified how organism size varies within species across space. Using a recently-digitised museum collection, we investigate at high intraspecific resolution how planktonic foraminifera, an important microfossil group, vary in size across the tropical and subtropical oceans of the world. We measured 3799 individuals of nine species in 53 seafloor sediments and analysed potential size biases in the museum collection. For each site, we obtained corresponding local values of mean annual sea-surface temperature (SST), net primary productivity (NPP), and relative abundance of each species. Given former studies, we expected species to reach largest shell sizes under optimal environmental conditions. At species level, we find the expected pattern; however, at population level, species greatly differ in how much size variation is explained by SST, NPP and/or relative abundance. While some species show a high predictability of size variation given one single variable *(Trilobatus sacculifer, Globigerinella siphonifera, Pulleniatina obliquiloculata, Globorotalia truncatulinoides),* other species showed either weak or no relationships between size and the studied covariates *(Globigerinoides ruber, G. conglobatus, Neogloboquadrina dutertrei, G. menardii, Globoconella inflata*). By incorporating intraspecific variation and sampling broader geographical ranges compared to previous studies, we conclude that shell size variation in planktonic foraminifera species cannot be consistently predicted by the environment. Our results caution against the general use of size as a proxy for planktonic foraminifera environmental optima, and emphasise the need for more studies analysing their within-species size variation. More generally, our findings show that contrasting results can be obtained when analysing patterns at different organisational levels, and highlights the utility of natural history collections.

## 1 Introduction

The size structure of plankton communities is an important determinant of the functions they realise in marine ecosystems (Barton et al., 2013; Litchman et al., 2013). However, most studies have focused on size distributions of assemblages (irrespective of species) or interspecific (among-species) variation instead of intraspecific (among-individual) variation (Sommer et al., 2017). Intraspecific variation can affect community dynamics (Des Roches et al., 2018), and influence species’ responses to environmental change (Mousing et al., 2017). Thus, by ignoring intraspecific variation, we have an incomplete understanding of the functions different plankton species perform in the ecosystem.

Planktonic foraminifera are an interesting group for studying intraspecific size variation. They are unicellular zooplankton that occur across the world’s oceans at low diversities (48 currently recognised species; Siccha and Kucera, 2017), and produce calcium carbonate tests (or shells). Upon death, their shells sink and accumulate on the ocean floor. The abundance of their shells preserved in marine sediments allows estimates of population-level size variation on a global scale (e.g., Schmidt et al. 2004a). Moreover, these abundant shells play a key role in the ocean carbon cycle (Schiebel, 2002) and, under good sedimentary conditions, yield a remarkably complete fossil record. This fossil record allows the quantification of how plankton shell size varied in geological time scales and responded to past environmental changes (Schmidt et al., 2004b). Thus, planktonic foraminifera can help elucidate the role of intraspecific variation in marine ecosystems across space and in deep time. However, a quantification of individual size variation across large biogeographical ranges is missing, limiting our understanding of what controls their within-species size variation.

Planktonic foraminifera grow in the pelagic zone by sequential addition of chambers until reproduction, when the gametes are released (gametogenesis), the cell dies and the empty shell sinks to the ocean floor (Be, 1976; Hemleben et al., 1989). Planktonic foraminifera assemblages show a global two-fold increase in average size from the poles to the tropics, and this increase correlates strongly and positively with sea surface temperature (Schmidt et al., 2004a). This pattern is opposite to the expected under the metabolic theory, in which higher temperatures affect metabolic rates positively, resulting in faster generation times and smaller organisms in the tropics (Brown et al., 2004; Savage et al., 2004). Planktonic foraminifera generation times seem to be constrained by their synchronous sexual reproduction (Bijma et al., 1990; Bijma and Hemleben, 1994; Jonkers et al., 2015; Venancio et al., 2016), and the final shell size likely determines the number of gametes released during gametogenesis (Hemleben et al., 1989). Larger sizes have been often associated to higher reproductive success in planktonic foraminifera (e.g., Grigoratou et a. 2019), although a quantification of this relationship is still absent as their reproduction has not been observed in the laboratory.

Previous studies have looked at planktonic foraminifera biogeographical size distribution and found that maximum shell size often coincides with maximum relative abundance, and occurs at specific optimum temperatures (optimumsize hypothesis; Kennett 1976; Hecht 1976; Malmgren and Kennett 1976, 1977; Kahn 1981; Schmidt et al. 2004a; Moller et al. 2013; see Be et al. 1973 for an exception). The local abundance of planktonic foraminifera species is usually estimated by counting dead assemblages from ocean floor sediments (Siccha and Kucera, 2017). This methodology often yields species’ relative abundances (relative to the co-occurring species in the sample) instead of absolute abundances.

Yet assemblages from the ocean floor sediments have the advantage of averaging out short-term fluctuations that might blur macroecological patterns (Kidwell and Tomasovych, 2013). Most of the studies supporting the optimum-size hypothesis focused on sediment samples collected within a single oceanic basin (Be et al., 1973; Kennett, 1976; Hecht, 1976; Malmgren and Kennett, 1976, 1977; Kahn, 1981; Moller et al., 2013), and thus a limited part of each species’ biogeographical range. The exception is the global study of Schmidt et al. (2004a), who analysed 69 Holocene samples worldwide. Although Schmidt et al. (2004a) were concerned with size variation of whole assemblages instead of within-species variation, they showed a strong 1:1 relationship between the temperatures at which a species reaches its largest size and its highest relative abundance, supporting the idea that planktonic foraminifera species reach largest shell sizes at their environmental optima. However, Schmidt et al. (2004a) only taxonomically identified a small fraction of their measured individuals. Thus, we lack studies testing the intraspecific consistency of the optimum-size hypothesis. If species reach largest shell sizes at their environmental optima, this pattern should be evident among their populations (within species).

Nutrient availability can mediate the temperature-size relationships observed in the plankton (Peter and Sommer, 2013; Marañón, 2015) and has been shown experimentally to affect planktonic foraminifera size: more food facilitates faster cell growth and larger final shell size (Be et al., 1981; Takagi et al., 2018). Planktonic foraminifera are omnivorous, preying on other plankton including diatoms, dinoflagellates, ciliates and copepods (Anderson et al., 1979; Spindler et al., 1984; Schiebel and Hemleben, 2017). Thus, phytoplankton are part of the foraminiferal diet and also attract other zooplankton predated by them (Northcote and Neil, 2005). If net primary productivity can be used as a proxy for planktonic foraminifera food availability, we expect size to correlate positively with primary productivity. Yet some species of planktonic foraminifera are photosymbiotic with eukaryotic algae (Hemleben et al., 1989; Takagi et al., 2019). Although photosymbiosis can not be the only form of daily nutrition to the foraminifer (Takagi et al., 2018), these photosymbiotic species usually occur in tropical-subtropical oligotrophic waters, suggesting photosymbiosis is an ecological strategy to survive in nutrient-limited environments (Be and Hutson, 1977). The expected relationship between net primary productivity and planktonic foraminifera size is not clear and has never been tested on a macroecological scale and high intraspecific resolution.

Here, we quantify the relationship between planktonic foraminifera within-species size variation and mean annual sea surface temperature (SST), mean annual net primary productivity (NPP) and relative abundance, plus the interaction between SST and NPP. We built a new population-level shell size dataset for nine extant species including the Pacific, Indian and Atlantic tropical and subtropical oceans. We expect size to ***(i)*** increase with increasing SST for tropical species and ***(ii)*** reach largest values at intermediate SST for transitional species (Schmidt et al., 2006). We also expect ***(iii)*** species to reach larger sizes where there are more resources available (i.e., higher NPP). As SST and NPP correlate in the open ocean (Schmidt et al., 2006), these two variables might interact and jointly predict more of the observed size variation than when tested alone. The optimum-size hypothesis also predicts a ***(iv)*** positive relationship between local relative abundance and size (Hecht, 1976), and ***(v)*** a strong, positive relationship between the SST at which each species reaches its largest size and its maximum abundance (Schmidt et al., 2004a).

## 2. Material and Methods

Our size dataset was extracted from the Henry Buckley Collection of Planktonic Foraminifera, held at the Natural History Museum in London, UK (NHMUK) (Rillo et al., 2016). We measured shell area of 3799 individuals from the nine species most commonly represented in the collection across 53 sites world-wide (Fig. 1). For each sampled site, we obtained corresponding data on the mean annual values of SST, NPP and relative abundance of each species. All the data visualisation and analyses were performed in the R programming language (version 3.3.3, R Core Team 2017).

**Figure 1:**
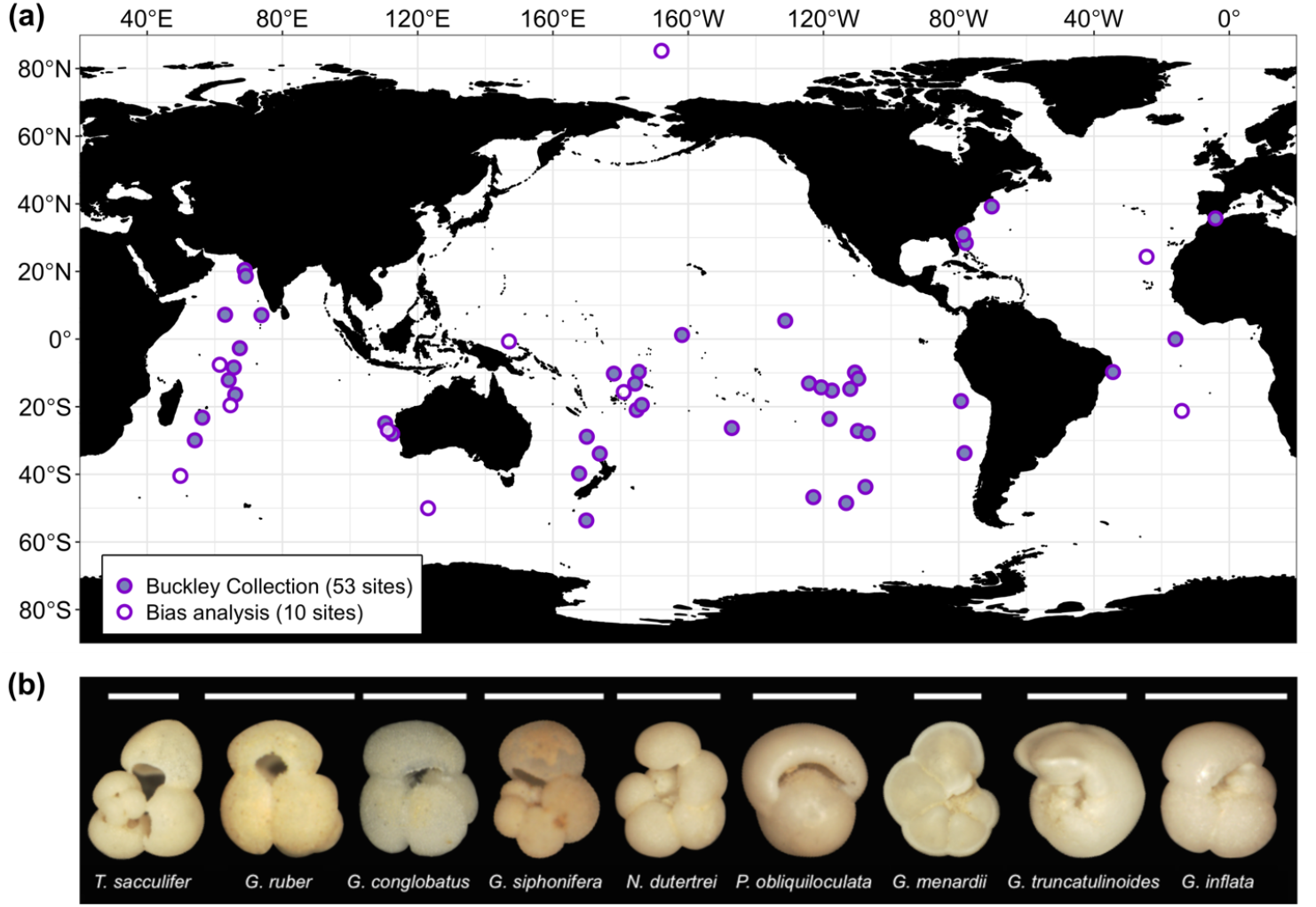
(a) The geographic distribution of the samples used from the Henry Buckley Collection (see text). Each dot on the map includes data on planktonic foraminifera shell size distributions, as well as corresponding data on mean annual values of sea surface temperature, net primary productivity and relative abundance. The purple dots represent the ten samples that were used to analyse the biases in the museum collection. The sample above 80°N was used only in the collection bias analysis. (b) A representative specimen from the collection for each species analysed. White bars represent 0.5 mm. From left to right: *Trilobatus sacculifer*, *Globigerinoides ruber*, *Globigerinoides conglobatus, Globigerinella siphonifera*, *Neogloboquadrina dutertrei, Pulleniatina obliquiloculata, Globorotalia menardii, Globorotalia truncatulinoides* and *Globoconella inflata.*

### 2.1 Study sites and samples

To amass the Henry Buckley Collection of Planktonic Foraminifera, Henry Buckley sampled 122 marine sediments from the NHMUK Ocean-Bottom Deposits Collection (OBD), which were collected by historical marine expeditions between 1873 and 1965 (Table S1) (Rillo et al., 2016). Sample processing usually consists of washing the sediment and dry sieving it over a 150μm sieve, then sampling the coarser fraction for planktonic foraminifera (Al-Sabouni et al. 2007). From the 122 samples processed by Buckley, we selected those that contained only extant species within the upper 15 cm of sediment and included at least one of the nine studied species (Fig. 1b). This resulted in 53 study sites predominantly in the tropical and subtropical regions of the Pacific, Indian and Atlantic oceans (Fig. 1a).

We determined the water depth for each site by matching the collection’s reported latitudes and longitudes to the ETOPO1 database hosted at the National Oceanic and Atmospheric Administration website (Amante and Eakins, 2009) using a 2 arc-minute grid resolution (R package *marmap* version 0.9.5; Pante and Simon-Bouhet 2013). Water depth ranged from 746 to 5153 meters below sea level (median 3296 m). Ten of the 53 samples in our dataset come from sediments prone to dissolution (i.e., waters deeper than 4000 meters for newly sedimented shells; Berger and Piper 1972). Dissolution may affect species size distributions, as smaller individuals are more prone to dissolution (Kennett, 1976), but we found no evidence that water depth is related to the size variation observed in our data (Table S2).

### 2.2 Shell size data

We measured cross-sectional shell area of the nine most abundant planktonic foraminifera species in the Buckley Collection. Each species has at least 244 specimens in the collection, resulting in 3799 individuals (Table 1, Fig. 2). The specimens were imaged using a Zeiss Axio Zoom V16 microscope and ZEN software at a resolution of 2.58 μm x 2.58 μm per pixel. Individual size was estimated based on the two-dimensional image of the specimen using a bespoke macro in Image-Pro Premier (version 9.1) that automatically recognises each specimen and measures its shell area. Henry Buckley mounted most specimens on the slides in a standard orientation (Fig. 1b, Table 1); individuals with a different orientation or dubious taxonomic identification were excluded from the analysis. Brombacher et al. (2017) quantified the reproducibility of shell area measurements and concluded that this two-dimensional size metric is highly consistent across slight deviations in mounting orientation. We avoided re-mounting the slides to preserve the Buckley Collection.

**Figure 2:**
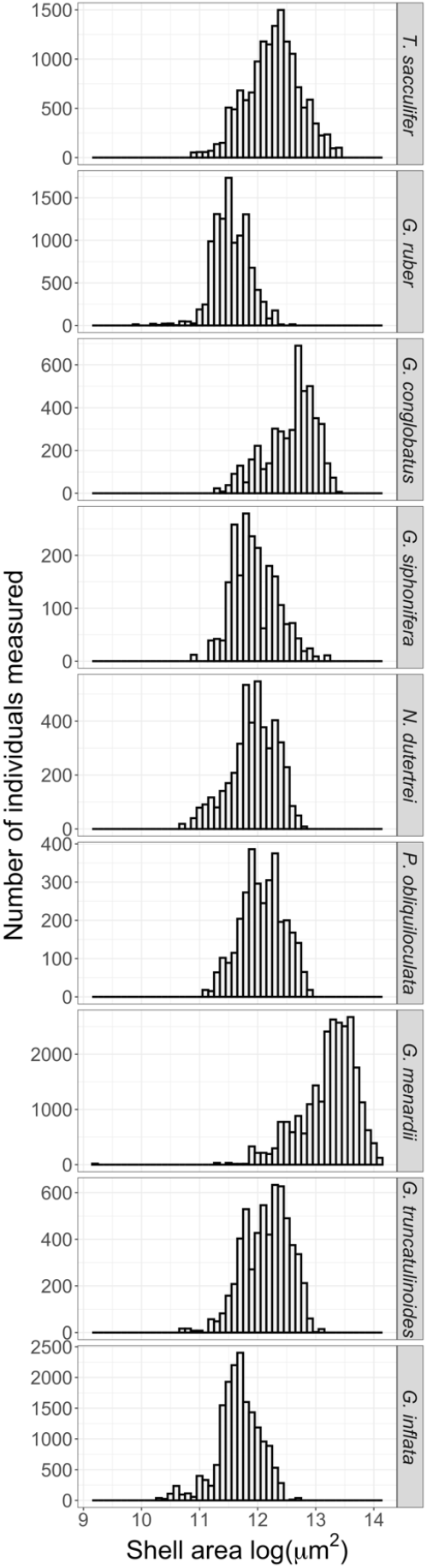
Shell size histograms (natural logarithm of the area) for each species of planktonic foraminifera present in the morphometric dataset. A total of 3799 individuals were measured.

**Table 1:**
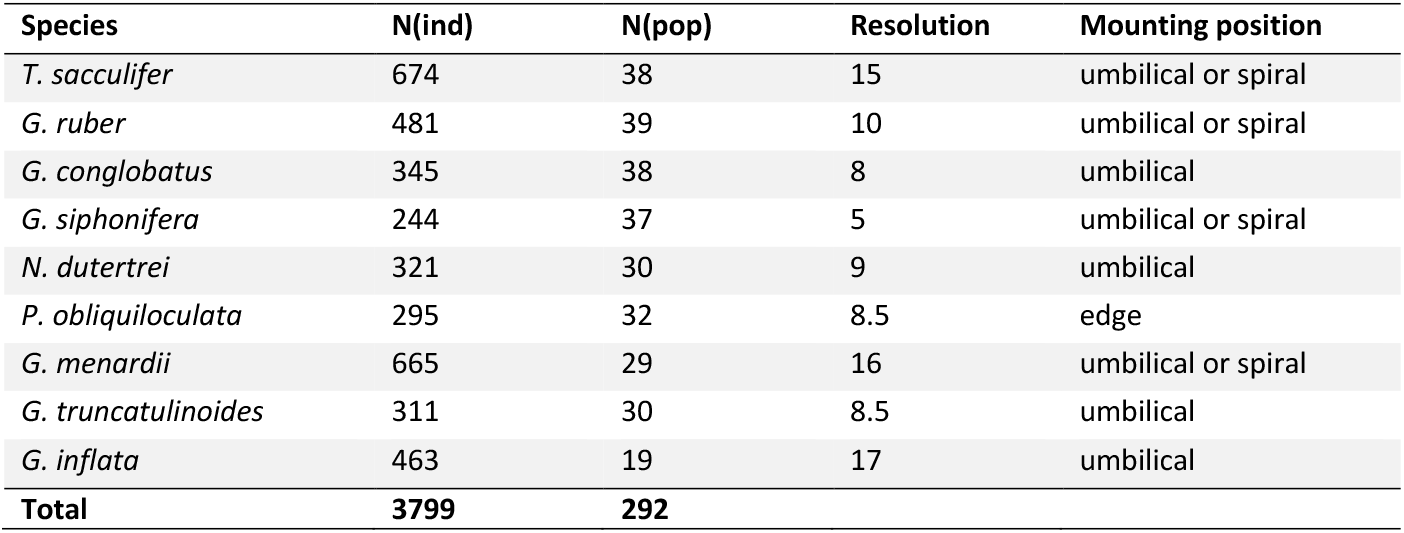
Overview of the morphometric dataset extracted from the museum collection. Columns: species names; number of individuals measured; number of populations per species (i.e., number of geographical sites per species); species resolution (i.e., median number of individuals per sample); mounting position in the collection (i.e., position in which the individuals of each species were measured).

The Buckley Collection could exhibit a collector effort bias, typically believed to occur towards larger specimens. To assess this potential bias, we re-sampled ten original bulk sediments from the OBD Collection that Buckley had used to amass his collection (filled dots in Fig. 1a, Table S3). We processed these ten samples (see Supplementary Information), and mounted species-specific slides to extract shell size data in the same way as for the original Buckley Collection (described above). We then compared the shell size distributions between the Buckley Collection samples and our samples. This comparison included 2873 individuals (1824 from the samples picked by us and 1049 from the Buckley Collection), across 65 populations from 20 species collected from the ten sites. We log-transformed the shell area data and calculated the mean, median, 75^th^ percentile, 95^th^ percentile and maximum value of each population shell size distribution. We then regressed each of these five population-metrics of the Buckley Collection against our re-sampled data (e.g., Fig. 3b), and calculated the residuals assuming 1:1 correspondence (i.e., the identity function). The residuals of the regressions are predominantly positive (Fig. 3a), indicating that the Buckley Collection has a consistent collector bias towards large specimens. As Henry Buckley personally carried out all the sample processing, isolation of foraminiferal specimens and their identification, the collector biases in his collection are likely to be systematic for within-species comparisons.

**Figure 3:**
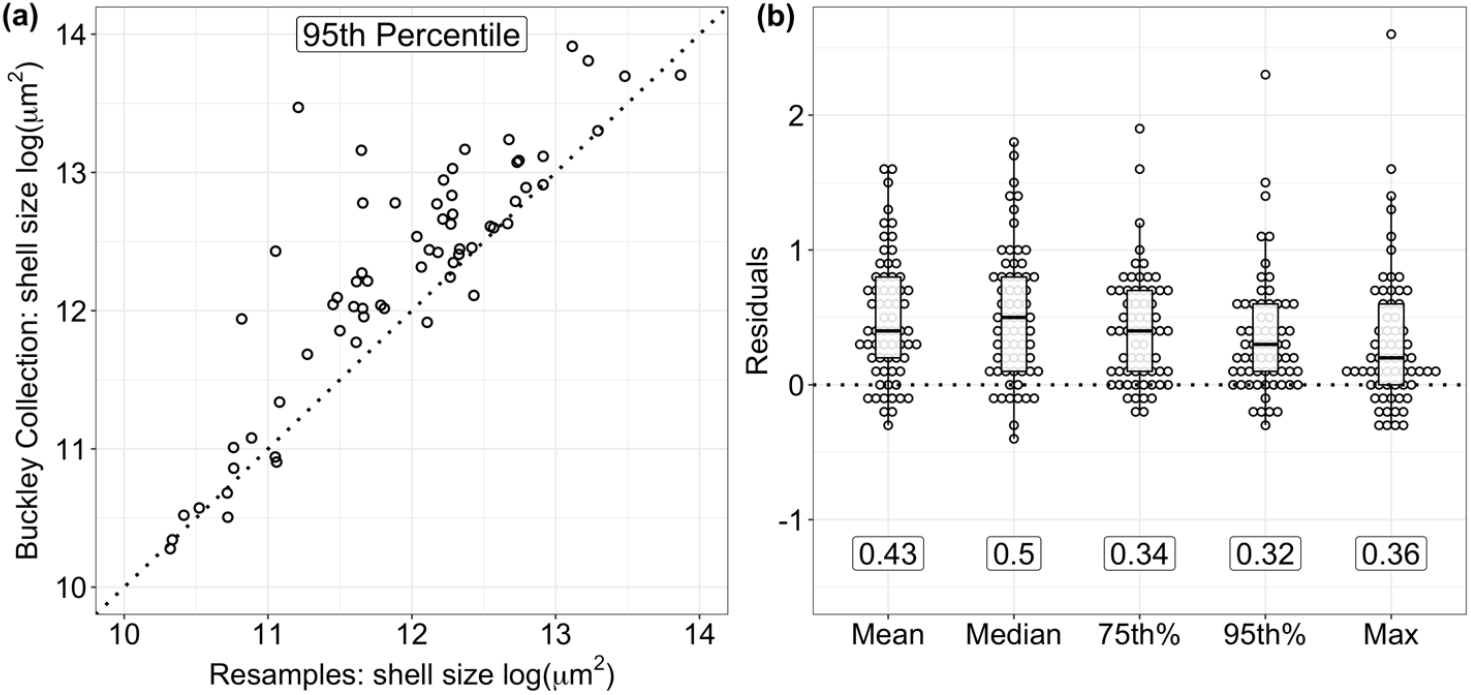
Bias analysis of the museum collection: difference in shell size distributions between data sampled by us and of the original Buckley Collection. **(a)** Plot of the population shell size distributions (95^th^ percentile) from the Buckley Collection against the re-sampled samples, dotted line represents the identity function (1:1 relationship). **(b)** Residuals were calculated between the Buckley Collection and our re-sampled samples with respect to the identity function (dotted line), using log-transformed population shell sizes. Numbers indicate mean squared error (MSE) of each metric. The 95^th^ percentile has the lowest MSE of the five metrics. See also Supplementary Information.

The mean squared error is lowest for the 95^th^ percentile (Fig. 3), meaning that this metric is the most representative population metric of the Buckley Collection. The robustness of the 95^th^ percentile of size distributions has also been documented by Schmidt et al. (2004a), as it is less sensitive to single outliers than the maximum value, and to representative sampling at the lower end of the size range than the mean and median values. Accordingly, we used the 95^th^ percentiles of the population shell size distributions as the dependent variable to investigate the covariates of planktonic foraminifera intraspecific shell size variation.

### 2.3 Sea-surface temperature data

We compiled mean annual values of SST from the World Ocean Atlas 2013 (WOA13, 0 meters depth, Locarnini et al. 2013) for each morphometric sample by matching its unique latitude and longitude coordinates to the nearest WOA13 1° grid point (approximately 111 km at the equator). The distances between the datasets were calculated using the World Geodetic System of 1984 (WGS 84) and R package *geosphere* (version 1.5-7; Hijmans 2015). We used SST data from the earliest decade available in the WOA13 database, resulting in SST data averaged for the years between 1955 and 1964. We chose this time period because the latest historical expedition associated with our morphometric dataset sailed in 1965 (Table S1).

### 2.4 Net-primary productivity data

We compiled mean annual values of NPP from the Ocean Productivity website (http://www.science.oregonstate.edu/ocean.productivity/). We selected the SeaWiFS estimates (based on the Eppley variation of the VGPM algorithm; Behrenfeld and Falkowski 1997) because they provide the earliest NPP data (starting in late 1997). We matched each morphometric sample coordinate to its nearest NPP sample as described for SST. The median distance between the datasets was 15 km. We considered only full years of NPP data collection, from January 1998 until December 2007.

### 2.5 Relative abundance data

To test for the relationship between population shell size and abundance (Hecht, 1976; Schmidt et al., 2004a), we extracted assemblage composition data from the ForCenS database (Siccha and Kucera, 2017). This database is a synthesis of planktonic foraminifera assemblage counts from surface sediment samples with 4205 records from unique sites worldwide, each with corresponding information on species’ relative abundance. We retrieved species relative abundance data for each morphometric sample by matching its coordinates to its nearest neighbour in the ForCenS database as described for SST. The median distance between the datasets was 106 km.

The spatial arrangement of shells on the seafloor is affected during settling by subsurface currents (Berger and Piper, 1972). Recent models estimate that settling foraminiferal shells can travel a maximum distance of 300 km in regions with large horizontal velocities (e.g., along the equator, in the western boundary currents and in the Southern Ocean; Van Sebille et al. 2015). To account for this post-mortem spatial variation of foraminiferal abundance on the seafloor, we retrieved ForCenS abundance data within a 300 km radius distance of each morphometric sample coordinate, which would be the maximum error according to Van Sebille et al. (2015). We then calculated the median relative abundance of each species based on all ForCenS samples that fell within 300 km of the morphometric sample. The analysis considering all ForCenS samples within 300 km produced consistent results compared to that using the nearest ForCenS sample (Table S5). Thus, we discuss results given the single nearest ForCenS sample.

### 2.6 Statistical analysis

For each species, the dependent response variable was the log-transformed size distribution (95^th^ percentile of each population) whereas the independent explanatory variables were the local mean annual SST, NPP and relative abundance. Linear and quadratic relationships were considered between shell size and SST and NPP, as well as the interaction between SST and NPP. Model fit and selection was assessed using Akaike information criterion corrected for small sample size (AICc); models within a difference of two AICc units (i.e., ΔAICc < 2) are equally plausible. Adjusted R squared (R_adj_^2^) were calculated for each model using the R package *rsq* (version 1.0.1; Zhang, 2017). Visual inspection of the residual plots did not reveal any obvious deviations from homoscedasticity, except for *G. inflata* (Fig. S1).

We used the observations in the 53 samples to investigate the optimum-size pattern at species-level. We calculated the SST at which each species reaches its largest size (Buckley Collection) and the SST at which the species is most abundant (ForCenS database). We tested the model of a 1:1 relationship between SST at largest size and at highest abundance, as these SST values are expected to be the same under the optimum-size hypothesis (Schmidt et al., 2004a).

## 3 Results

Size variation within species of planktonic foraminifera is high (Fig. 2, 4) and can range over one order of magnitude among adults of the same species (e.g., from 150*μ*m to 1500*μ*m *Globorotalia menardii,* Fig. 2). This high intraspecific variation relates differently to SST, NPP and relative abundance for the different species. When tested alone, SST shows a significant relationship with size in four out of nine species. *T. sacculifer, G. siphonifera* and *P. obliquiloculata* increase in size significantly with a linear increase of SST, whereas *G. truncatulinoides* size variation is significantly explained by a quadratic function of SST (Fig. 4a). Shell sizes in the other five species do not co-vary significantly with SST. NPP and size variation show a significant positive relationship in *G. siphonifera* and a significant negative relationship in *G. truncatulinoides* (Fig. 4b). Local relative abundance explains less size variation than SST and NPP for all species except *G. menardii* and *G. inflata* (Fig. 4c). *T. sacculifer* shows a positive relationship between shell size and relative abundance, but weaker than with SST.

**Figure 4:**
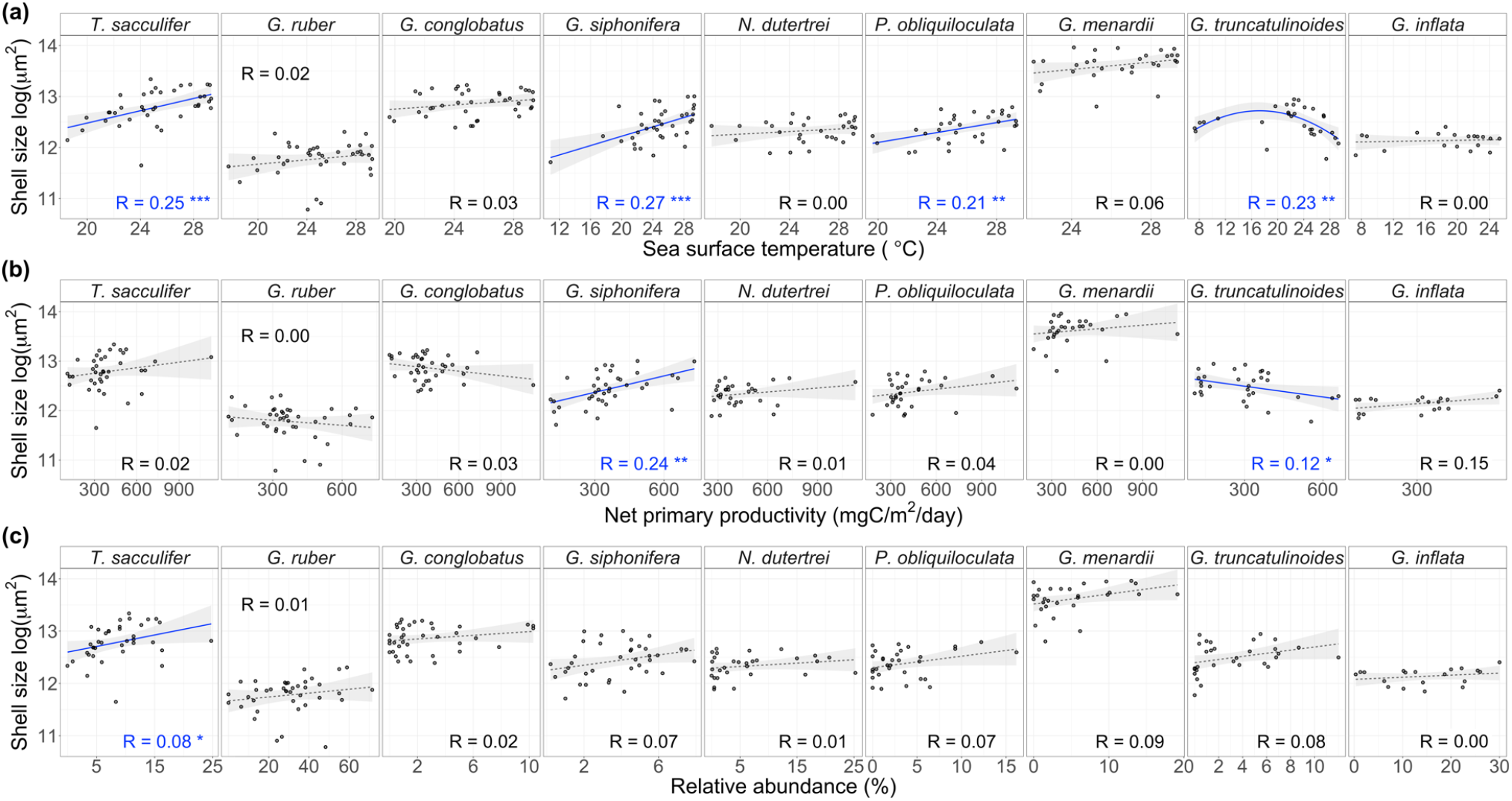
Natural logarithm of shell size (represented by the 95^th^ percentile of the population distribution) as a function of **(a)** mean annual sea-surface temperature, **(b)** mean annual net-primary productivity and **(c)** species’ local relative abundance. Solid blue lines show significant relationship whereas dashed grey lines non-significant; grey shades show standard error of the model. Most models are linear, except for the quadratic relationship between *G. truncatulinoides* size and temperature. The legend in each plot shows the adjusted R squared for each species. Significance codes: *** p<0.001; ** p<0.01; * p<0.05.

SST and NPP correlate (*corr* = 0.4, *p-value* < 0.001), thus their isolated effect on size might be confounding (Fig. 4a,b). For this reason, we considered all variables together in a model selection framework, including the interaction between SST and NPP. The explanatory power of SST, NPP and relative abundance still varies greatly among species (Table 2). SST explains 25% and 21% of the intraspecific size variation of *T. sacculifer* and *P. obliquiloculata,* respectively. SST and NPP together reach the highest predictability of size variation: 33% in *G. siphonifera* and 32% in *G. truncatulinoides.* When considering the interaction between SST and NPP, 34% of the size variation in *G. truncatulinoides* and 19% in *G. conglobatus* is explained. NPP alone explains 15% of the size variation of *G. inflata,* although the null model is equally plausible (ΔAICc < 2). *G. menardii* size variation can be predicted by 15% given the additive effect of SST and relative abundance, but the null model is also plausible. Surprisingly, *G. ruber* and *N. dutertrei* size variations are poorly predicted by any of the three variables (see also Fig. 4), being best explained by the null model (i.e., the sample mean).

**Table 2:**
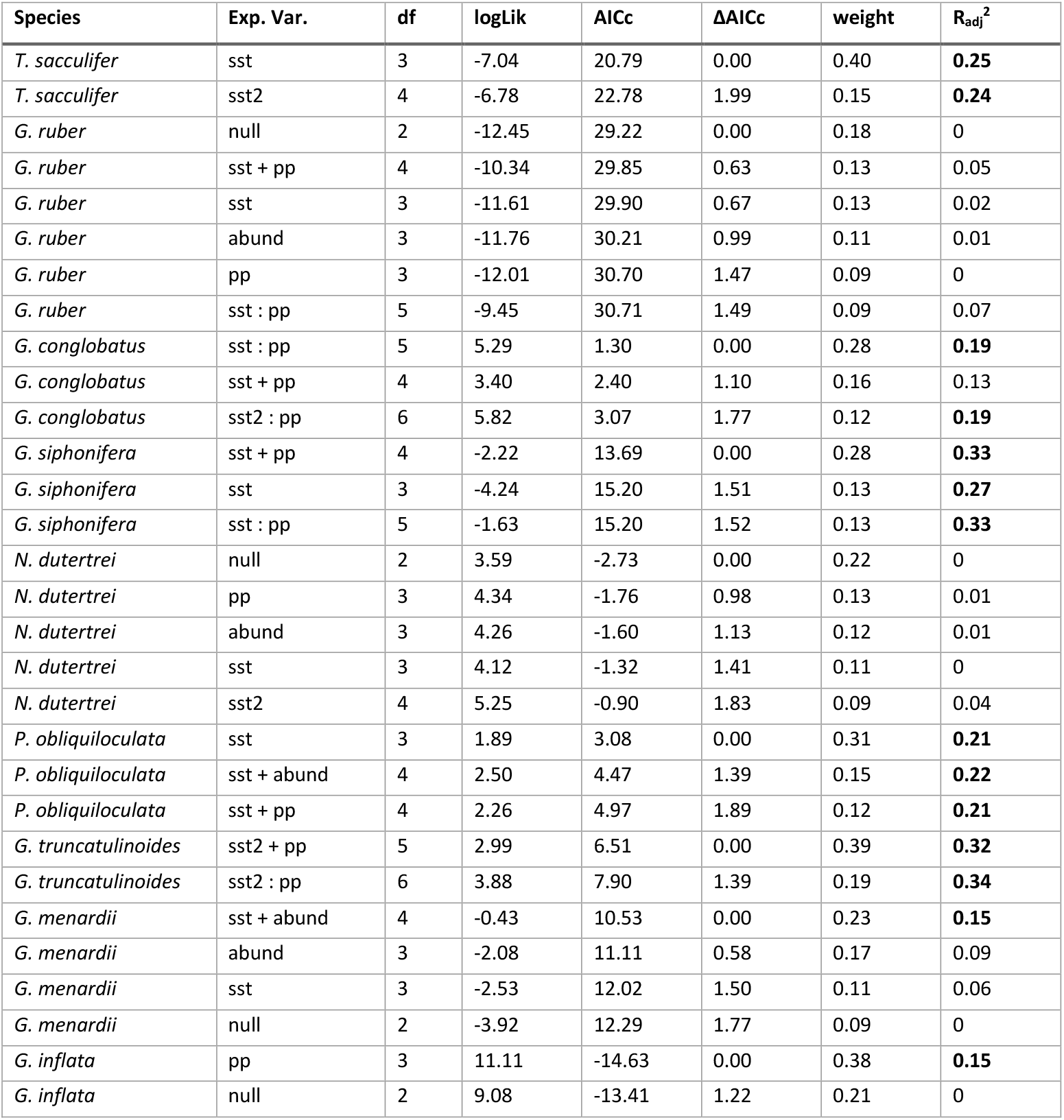
Model selection of the linear and quadratic models testing if planktonic foraminifera shell size (represented by the 95^th^ percentile of each population size distribution) can be predicted by mean annual sea-surface temperature (sst linear effect, sst^2^ quadratic effect), mean annual net-primary productivity (npp) and/or species’ relative abundance (abund). A model including the interaction between sst and npp (sst: npp) was also considered. Columns: species, explanatory variables, degrees of freedom, log-likelihood, Akaike Information Criterion corrected for small sample size (AICc), AICc difference between models (ΔAICc), model weight, adjusted R squared (bold values: above 0.15). All models within two ΔAICc units are shown and considered equally plausible.

| Species | Exp. Var. | df | logLik | AICc | ΔAICc | weight | $R_{adj}^2$ |
| --- | --- | --- | --- | --- | --- | --- | --- |
| *T. sacculifer* | sst | 3 | -7.04 | 20.79 | 0.00 | 0.40 | **0.25** |
| *T. sacculifer* | sst2 | 4 | -6.78 | 22.78 | 1.99 | 0.15 | **0.24** |
| *G. ruber* | null | 2 | -12.45 | 29.22 | 0.00 | 0.18 | 0 |
| *G. ruber* | sst + pp | 4 | -10.34 | 29.85 | 0.63 | 0.13 | 0.05 |
| *G. ruber* | sst | 3 | -11.61 | 29.90 | 0.67 | 0.13 | 0.02 |
| *G. ruber* | abund | 3 | -11.76 | 30.21 | 0.99 | 0.11 | 0.01 |
| *G. ruber* | pp | 3 | -12.01 | 30.70 | 1.47 | 0.09 | 0 |
| *G. ruber* | sst : pp | 5 | -9.45 | 30.71 | 1.49 | 0.09 | 0.07 |
| *G. globobatus* | sst : pp | 5 | 5.29 | 1.30 | 0.00 | 0.28 | **0.19** |
| *G. globobatus* | sst + pp | 4 | 3.40 | 2.40 | 1.10 | 0.16 | 0.13 |
| *G. globobatus* | sst2 : pp | 6 | 5.82 | 3.07 | 1.77 | 0.12 | **0.19** |
| *G. siphonifera* | sst + pp | 4 | -2.22 | 13.69 | 0.00 | 0.28 | **0.33** |
| *G. siphonifera* | sst | 3 | -4.24 | 15.20 | 1.51 | 0.13 | **0.27** |
| *G. siphonifera* | sst : pp | 5 | -1.63 | 15.20 | 1.52 | 0.13 | **0.33** |
| *N. dutertrei* | null | 2 | 3.59 | -2.73 | 0.00 | 0.22 | 0 |
| *N. dutertrei* | pp | 3 | 4.34 | -1.76 | 0.98 | 0.13 | 0.01 |
| *N. dutertrei* | abund | 3 | 4.26 | -1.60 | 1.13 | 0.12 | 0.01 |
| *N. dutertrei* | sst | 3 | 4.12 | -1.32 | 1.41 | 0.11 | 0 |
| *N. dutertrei* | sst2 | 4 | 5.25 | -0.90 | 1.83 | 0.09 | 0.04 |
| *P. obliquiloculata* | sst | 3 | 1.89 | 3.08 | 0.00 | 0.31 | **0.21** |
| *P. obliquiloculata* | sst + abund | 4 | 2.50 | 4.47 | 1.39 | 0.15 | **0.22** |
| *P. obliquiloculata* | sst + pp | 4 | 2.26 | 4.97 | 1.89 | 0.12 | **0.21** |
| *G. truncatulinoides* | sst2 + pp | 5 | 2.99 | 6.51 | 0.00 | 0.39 | **0.32** |
| *G. truncatulinoides* | sst2 : pp | 6 | 3.88 | 7.90 | 1.39 | 0.19 | **0.34** |
| *G. menardii* | sst + abund | 4 | -0.43 | 10.53 | 0.00 | 0.23 | **0.15** |
| *G. menardii* | abund | 3 | -2.08 | 11.11 | 0.58 | 0.17 | 0.09 |
| *G. menardii* | sst | 3 | -2.53 | 12.02 | 1.50 | 0.11 | 0.06 |
| *G. menardii* | null | 2 | -3.92 | 12.29 | 1.77 | 0.09 | 0 |
| *G. inflata* | pp | 3 | 11.11 | -14.63 | 0.00 | 0.38 | **0.15** |
| *G. inflata* | null | 2 | 9.08 | -13.41 | 1.22 | 0.21 | 0 |

Regarding species-level patterns, we tested the relationship between the SST where each species reaches its largest size and the SST where the species is most abundant (Fig. 5). Our data shows a positive, significant correlation (Fig. 5b) similarly to the findings of Schmidt et al. (2004a) (Fig. 5a).

**Figure 5:**
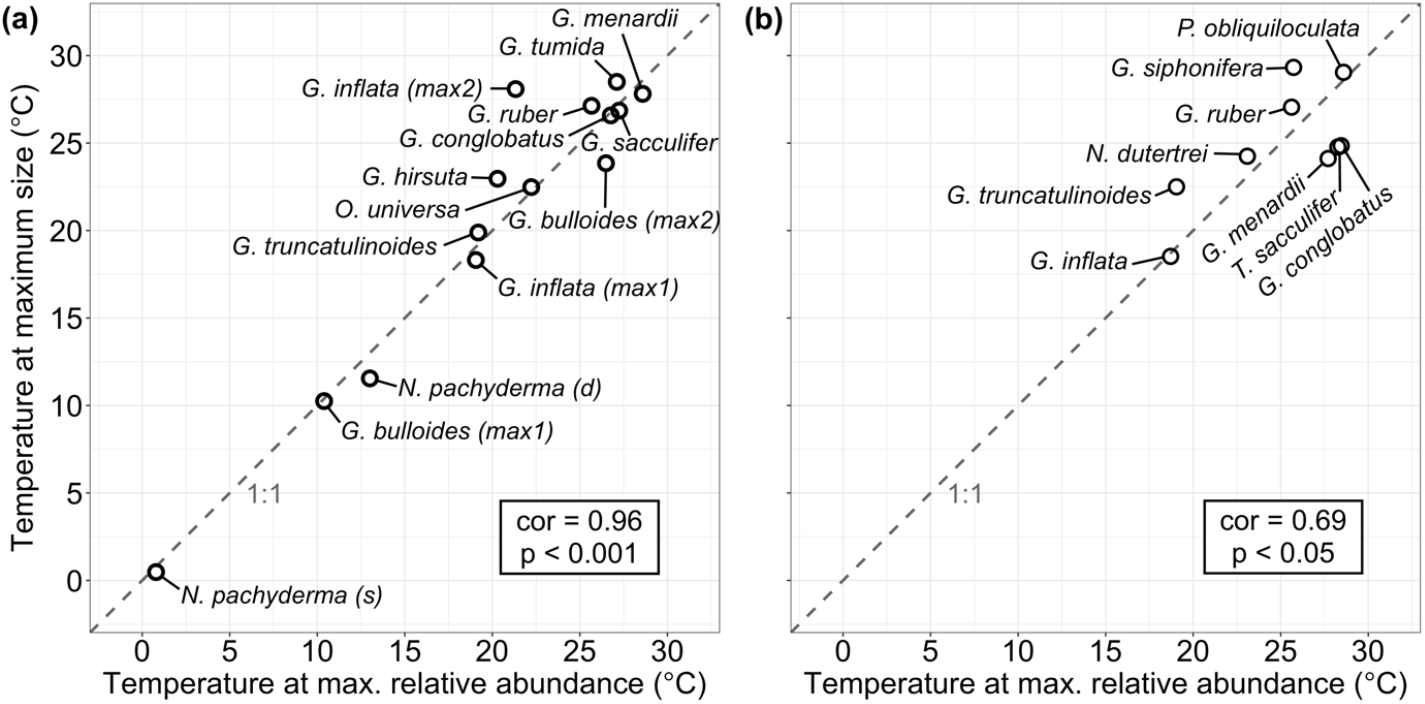
Sea surface temperatures where planktonic foraminifera species reach maximum shell size and maximum relative abundance in the surface sediments from **(a)** Schmidt et al. (2004a) and **(b)** this study. Dashed lines indicate 1:1 relationship.

## 4. Discussion

Our new morphometric dataset of species-resolved planktonic foraminifera allowed us to explore the correlates of intraspecific shell size variation at a macroecological scale. We expected planktonic foraminifera species to be larger at their environmental optima, identified here using sea surface temperature, net primary productivity and species’ relative abundances. At species level, the expected pattern is observed (Fig. 5); however, at population level (within-species), species differed in how much size variation is explained by these variables (Fig. 4, Table 2).

When tested alone, SST explained most of the shell size variation within planktonic foraminifera species, but only four of the nine studied species showed a significant relationship between size and SST (Fig. 4a). The tropical species *T. sacculifer, G. siphonifera* and *P. obliquiloculata* showed the expected positive linear relationship between SST and shell size, while the transitional *G. truncatulinoides* showed a quadratic relationship between shell size and SST (Fig. 4a). Results for these four species support previous observations that that planktonic foraminifera species are largest at their thermal optima (Schmidt et al., 2004a). However, the remaining five species (namely *G. ruber, G. conglobatus, G. menardii, N. dutertrei* and *G. inflata)* showed neither a linear or quadratic relationship between shell size and SST.

We found weak and conflicting evidence for links between shell size, open-ocean NPP and species’ symbiotic strategy. Six of the nine species studied are obligatory (persistent) photosymbiotic species (*G. siphonifera, T. sacculifer, G. ruber, G. conglobatus, N. dutertrei* and *G. menardii;* Hemleben et al., 1989; Takagi et al., 2019). Of these, only *G.siphonifera* and *G. conglobatus* size variation was best explained by a model that includes NPP (and SST, Table 2), but the relationship between size and NPP differed in sign and significance for these two (Fig. 4b). The facultative symbiotic species *P. obliquiloculata* and *G. inflata* (Takagi et al., 2019) showed non-significant relationships between size and NPP. And *G. truncatulinoides,* the only asymbiotic species, included NPP in all the best models explaining its intraspecific size variation (Table 2), but the relationship between size and NPP is negative, contrary to the expected if species would grow more when more nutrients are available. More generally, NPP was not a good predictor of size within the (mostly photosymbiotic) species studied here.

The presence of cryptic genetic diversity could have contributed to the high intraspecific variation found in our study. Some planktonic foraminifera species are in fact complexes of lineages, which are genetically independent but morphologically similar (Morard et al., 2015). Cryptic species have been shown to occupy different niches and be endemic to particular ocean basins (De Vargas et al., 1999; Darling and Wade, 2008; Weiner et al., 2014). So, the macroecological scale of our study likely includes cryptic diversity within our morphologically-defined species. *T. sacculifer* and *G. conglobatus* are the only genetically homogeneous species in our study (Aurahs et al., 2011; Seears et al., 2012; Andre et al., 2013). The other species have been shown to comprise some level of cryptic diversity (De Vargas et al., 1999; Darling and Wade, 2008; Morard et al., 2011; Aurahs et al., 2011; Ujiie et al., 2012; Quillevere et al., 2011; Weiner et al., 2014, 2015), except for *G. menardii,* whose genetic diversity has not yet been determined (Seears et al., 2012). The predictability of size variation in our study does not seem to relate directly to the cryptic diversity of the species: species with strong relationships between size and SST and/or NPP include not only the genetically homogeneous *T. sacculifer* and *G. conglobatus,* but also the genetically diverse *G. siphonifera, P. obliquiloculata* and *G. truncatulinoides.* Thus, our result suggests that shell sizes of these distinct genetic types likely respond to SST and/or NPP in similar ways.

The Buckley Collection contains samples from historical expeditions that collected marine sediments using devices such as a dredge, which potentially disturb the ocean floor surface and can recover a mix of Holocene (surface) and deeper, Pleistocene sediments (Rillo et al., 2019). Although this source of bias is inherent to this museum collection, as it includes samples from pioneering marine expeditions such as HMS *Challenger,* it can potentially increase the size variation observed and, consequently, blur biogeographical patterns because of the temporal mixing of samples. To assess this bias, we removed six samples recovered using dredges or grabers (Table S1) and re-ran analyses to test if different patterns emerge. Most species’ size patterns were unchanged except for *G. conglobatus,* whose best models without dredged samples include the null model, and *G. inflata,* whose best model includes only NPP (and not the null model as before) (Table S6). Overall the general pattern remains: intraspecific size variation in planktonic foraminifera cannot be explained consistently for all species.

The local relative abundance of a species was in general a poor predictor of its size variation (Table 2, Fig. 4c), contrary to the idea that planktonic foraminifera species are largest where they are most common (Hecht, 1976). The weak relationship between size and local relative abundance found in our study could be due to the fact that we used relative abundance data from a different source (the ForCenS database; Siccha and Kucera 2017). We assessed the robustness of our results by testing the optimum-size hypothesis on a more uniform, but smaller, dataset: the ten resampled bulk samples used for the bias analysis (Fig. 1a, 2; Table S3). We measured shell sizes and calculated species’ relative abundances for each of the ten assemblages, so that the same specimens were used to extract abundance and size data. 65 populations of 20 species were then used to test if population shell size could be predicted by relative abundance using a linear-mixed effect model with species as random effects. The results showed no significant relationship between size variation and relative abundance (chi-square test, χ^2^ = 2.18, P value = 0.14, Table S4), supporting our previous findings using the larger Buckley Collection data (Table 2, Fig. 4c).

Potentially the most significant source of uncertainty in the evaluation of the existence of a relationship between size and optimum environmental conditions is the use of relative, instead of absolute, abundance as an indicator of planktonic foraminifera ecological and thermal optima (Hecht, 1976; Kucera, 2007). When a species is less abundant (or longer-lived) than other species in the local plankton community, its relative abundance in the sediment will be strongly influenced by the more abundant (or shorter-lived) co-occurring species. Moreover, a species can reach high relative abundance because it is better able to tolerate stress than other species. As a result of stress, the abundances of all species will be low, but the most tolerant species will reach high relative abundances. The high relative abundances of *G. ruber* together with the lack of a relationship between shell size and the studied variables (Table 2, Fig. 4) suggests that this species is able to tolerate stress and, therefore, reach high relative abundance in sub-optimal conditions (Be and Tolderlund, 1971; Schmuker and Schiebel, 2002). Preferably the environmental optimum of a species would be determined by comparing absolute abundances of populations of the same species independently of the abundances of the other species in the assemblage. Further investigation on the relationship between size and optimal environmental conditions within species should focus on absolute abundance estimates derived from plankton tows and sediment traps (e.g., Beer et al. 2010; Aldridge et al. 2012; Weinkauf et al. 2016), instead of relative abundances in the sediment.

The idea that planktonic foraminifera species are largest at their environmental optima is largely based on the strong relationship between the SST at which a species reaches maximum size and the SST at which it reaches maximum relative abundance found by Schmidt et al. (2004a, Fig. 5a). Although we find support for this relationship at species level (Fig. 5b), the relationship is mostly absent within species (i.e., at population level, Fig. 4a), showing that species are not consistently larger at their environmental optimum. Thus, contrasting results can be obtained when analysing patterns at different organisational levels. Further attempts to reconcile the intra- and interspecific processes underlying organism size variation are necessary to bridge discrepancies observed among local and macroecological patterns.

## 5. Conclusion

We tested the hypothesis that planktonic foraminifera species are largest under optimal environmental conditions (Hecht, 1976; Schmidt et al., 2004a), identified using local SST, NPP and relative abundance of the species. We found a mixed picture: while *T. sacculifer, G. siphonifera, P. obliquiloculata* and *G. truncatulinoides* reach larger sizes at specific environmental conditions, *G. ruber, G. conglobatus, N. dutertrei, G. menardii* and *G. inflata* do not show statistically significant relationships between size and the studied environmental and ecological variables. Thus, our results caution against the idea that all planktonic foraminifera species grow larger at their environmental optima. More standardised individual-level studies are necessary to disentangle the mechanisms underlying planktonic foraminifera biogeographical size distributions. More generally, our work highlights the utility of natural history collections and emphasises the importance of studying intraspecific variation when interpreting macroecological patterns.

## Acknowledgements

We gratefully acknowledge Daniela N. Schmidt, Isabel S. Fenton and Andy Purvis for insightful discussions about the project. We also thank Anieke Brombacher for patient assistance with the Image-Pro Premier software and, together with Mauro T. C. Sugawara and Phil Fenberg, for helpful comments in earlier versions of this manuscript.

## Authors contribution

MCR designed the research question, with input from all the authors. MCR imaged and measured the museum specimens, with input from GM. MCR and MK identified the specimens for the museum bias analysis. MCR, MK and THGE designed the statistical analysis. MCR performed the analysis, made the figures, and wrote the manuscript. All authors reviewed the manuscript.

## Data Availability Statement

The data and the R code used to produce this analysis are available from the NHM Data Portal: https://doi.org/10.5519/0056541. Specimens’ images can be found at https://doi.org/10.5519/0035055.

## Supplementary Information

**Table S1:**
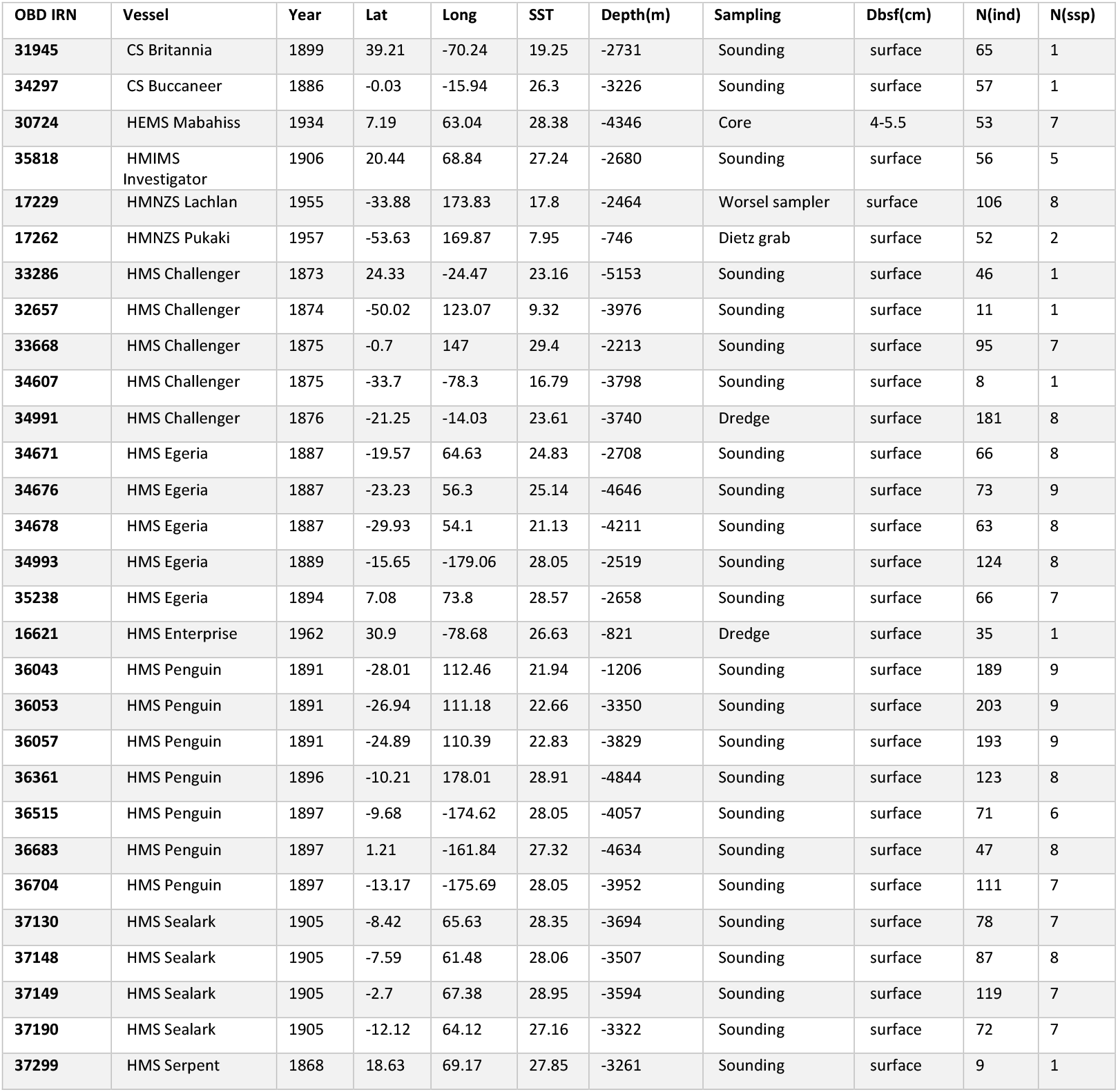
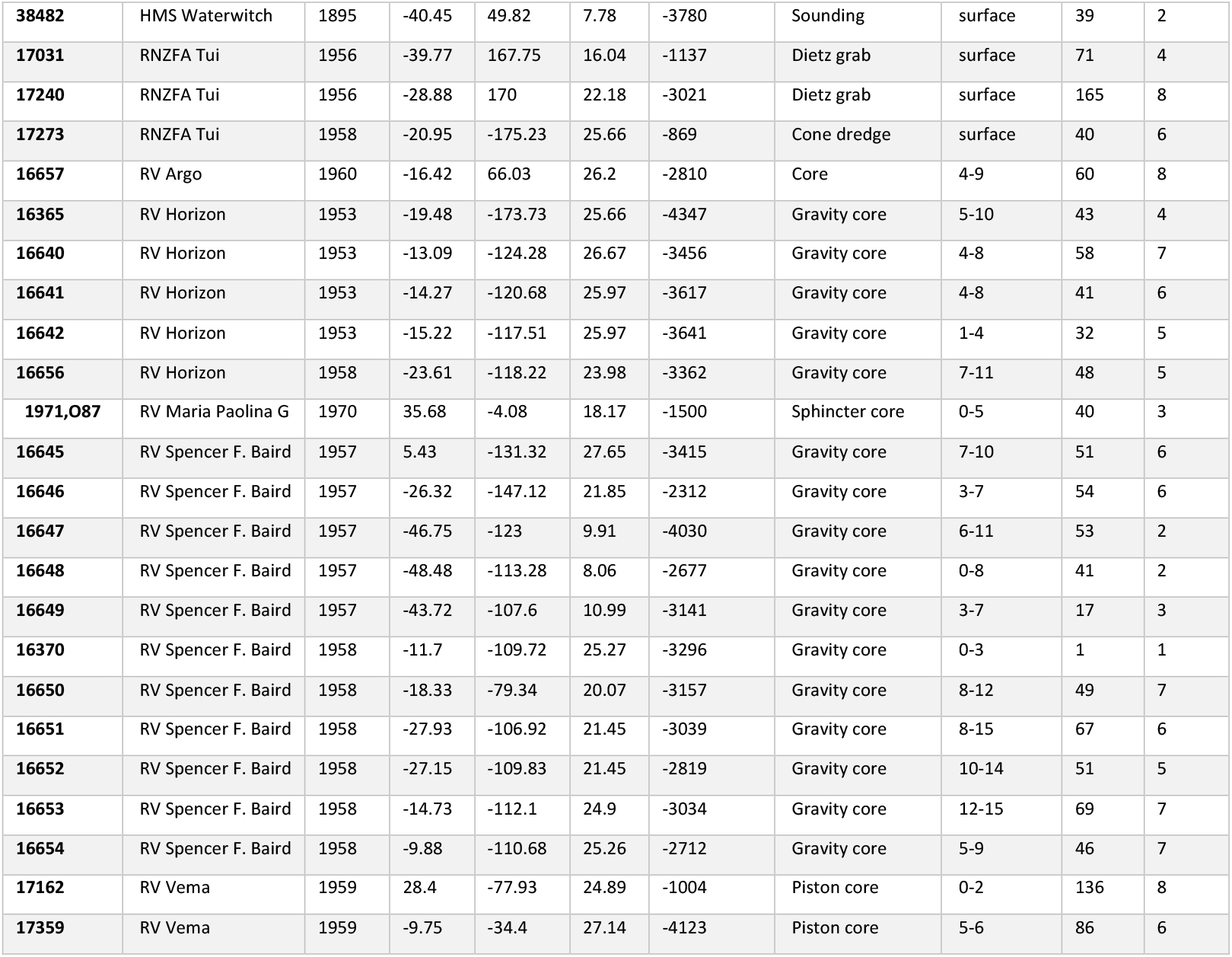
Samples from the Henry Buckley Collection of Planktonic Foraminifera at The Natural History Museum, London (NHMUK) used in the morphometric analysis. Columns: NHMUK Internal Record Number of the sediment in the Ocean-Bottom Deposits Collection (OBD IRN); name of the vessel that collected the sample; year the sample was collected; latitude (Lat) and longitude (Long) given in decimal degrees; sea surface temperature (SST) in Celsius degrees; water depth in meters; sampling method used in the historical expedition; depth below the seafloor (Dbsf) sampled in centimetres; number of planktonic Foraminifera specimens N(ind) and species N(ssp) measured at each site.

| OBD IRN | Vessel | Year | Lat | Long | SST | Depth(m) | Sampling | Dbsf(cm) | N(ind) | N(ssp) |
| --- | --- | --- | --- | --- | --- | --- | --- | --- | --- | --- |
| 31945 | CS Britannia | 1899 | 39.21 | -70.24 | 19.25 | -2731 | Sounding | surface | 65 | 1 |
| 34297 | CS Buccaneer | 1886 | -0.03 | -15.94 | 26.3 | -3226 | Sounding | surface | 57 | 1 |
| 30724 | HEMS Mabahiss | 1934 | 7.19 | 63.04 | 28.38 | -4346 | Core | 4-5.5 | 53 | 7 |
| 35818 | HMIMS Investigator | 1906 | 20.44 | 68.84 | 27.24 | -2680 | Sounding | surface | 56 | 5 |
| 17229 | HMNZS Lachlan | 1955 | -33.88 | 173.83 | 17.8 | -2464 | Worsel sampler | surface | 106 | 8 |
| 17262 | HMNZS Pukaki | 1957 | -53.63 | 169.87 | 7.95 | -746 | Dietz grab | surface | 52 | 2 |
| 33286 | HMS Challenger | 1873 | 24.33 | -24.47 | 23.16 | -5153 | Sounding | surface | 46 | 1 |
| 32657 | HMS Challenger | 1874 | -50.02 | 123.07 | 9.32 | -3976 | Sounding | surface | 11 | 1 |
| 33668 | HMS Challenger | 1875 | -0.7 | 147 | 29.4 | -2213 | Sounding | surface | 95 | 7 |
| 34607 | HMS Challenger | 1875 | -33.7 | -78.3 | 16.79 | -3798 | Sounding | surface | 8 | 1 |
| 34991 | HMS Challenger | 1876 | -21.25 | -14.03 | 23.61 | -3740 | Dredge | surface | 181 | 8 |
| 34671 | HMS Egeria | 1887 | -19.57 | 64.63 | 24.83 | -2708 | Sounding | surface | 66 | 8 |
| 34676 | HMS Egeria | 1887 | -23.23 | 56.3 | 25.14 | -4646 | Sounding | surface | 73 | 9 |
| 34678 | HMS Egeria | 1887 | -29.93 | 54.1 | 21.13 | -4211 | Sounding | surface | 63 | 8 |
| 34993 | HMS Egeria | 1889 | -15.65 | -179.06 | 28.05 | -2519 | Sounding | surface | 124 | 8 |
| 35238 | HMS Egeria | 1894 | 7.08 | 73.8 | 28.57 | -2658 | Sounding | surface | 66 | 7 |
| 16621 | HMS Enterprise | 1962 | 30.9 | -78.68 | 26.63 | -821 | Dredge | surface | 35 | 1 |
| 36043 | HMS Penguin | 1891 | -28.01 | 112.46 | 21.94 | -1206 | Sounding | surface | 189 | 9 |
| 36053 | HMS Penguin | 1891 | -26.94 | 111.18 | 22.66 | -3350 | Sounding | surface | 203 | 9 |
| 36057 | HMS Penguin | 1891 | -24.89 | 110.39 | 22.83 | -3829 | Sounding | surface | 193 | 9 |
| 36361 | HMS Penguin | 1896 | -10.21 | 178.01 | 28.91 | -4844 | Sounding | surface | 123 | 8 |
| 36515 | HMS Penguin | 1897 | -9.68 | -174.62 | 28.05 | -4057 | Sounding | surface | 71 | 6 |
| 36683 | HMS Penguin | 1897 | 1.21 | -161.84 | 27.32 | -4634 | Sounding | surface | 47 | 8 |
| 36704 | HMS Penguin | 1897 | -13.17 | -175.69 | 28.05 | -3952 | Sounding | surface | 111 | 7 |
| 37130 | HMS Sealark | 1905 | -8.42 | 65.63 | 28.35 | -3694 | Sounding | surface | 78 | 7 |
| 37148 | HMS Sealark | 1905 | -7.59 | 61.48 | 28.06 | -3507 | Sounding | surface | 87 | 8 |
| 37149 | HMS Sealark | 1905 | -2.7 | 67.38 | 28.95 | -3594 | Sounding | surface | 119 | 7 |
| 37190 | HMS Sealark | 1905 | -12.12 | 64.12 | 27.16 | -3322 | Sounding | surface | 72 | 7 |
| 37299 | HMS Serpent | 1868 | 18.63 | 69.17 | 27.85 | -3261 | Sounding | surface | 9 | 1 |
| 38482 | HMS Waterwitch | 1895 | -40.45 | 49.82 | 7.78 | -3780 | Sounding | surface | 39 | 2 |
| 17031 | RNZFA Tui | 1956 | -39.77 | 167.75 | 16.04 | -1137 | Dietz grab | surface | 71 | 4 |
| 17240 | RNZFA Tui | 1956 | -28.88 | 170 | 22.18 | -3021 | Dietz grab | surface | 165 | 8 |
| 17273 | RNZFA Tui | 1958 | -20.95 | -175.23 | 25.66 | -869 | Cone dredge | surface | 40 | 6 |
| 16657 | RV Argo | 1960 | -16.42 | 66.03 | 26.2 | -2810 | Core | 4-9 | 60 | 8 |
| 16365 | RV Horizon | 1953 | -19.48 | -173.73 | 25.66 | -4347 | Gravity core | 5-10 | 43 | 4 |
| 16640 | RV Horizon | 1953 | -13.09 | -124.28 | 26.67 | -3456 | Gravity core | 4-8 | 58 | 7 |
| 16641 | RV Horizon | 1953 | -14.27 | -120.68 | 25.97 | -3617 | Gravity core | 4-8 | 41 | 6 |
| 16642 | RV Horizon | 1953 | -15.22 | -117.51 | 25.97 | -3641 | Gravity core | 1-4 | 32 | 5 |
| 16656 | RV Horizon | 1958 | -23.61 | -118.22 | 23.98 | -3362 | Gravity core | 7-11 | 48 | 5 |
| 1971,087 | RV Maria Paolina G | 1970 | 35.68 | -4.08 | 18.17 | -1500 | Sphincter core | 0-5 | 40 | 3 |
| 16645 | RV Spencer F. Baird | 1957 | 5.43 | -131.32 | 27.65 | -3415 | Gravity core | 7-10 | 51 | 6 |
| 16646 | RV Spencer F. Baird | 1957 | -26.32 | -147.12 | 21.85 | -2312 | Gravity core | 3-7 | 54 | 6 |
| 16647 | RV Spencer F. Baird | 1957 | -46.75 | -123 | 9.91 | -4030 | Gravity core | 6-11 | 53 | 2 |
| 16648 | RV Spencer F. Baird | 1957 | -48.48 | -113.28 | 8.06 | -2677 | Gravity core | 0-8 | 41 | 2 |
| 16649 | RV Spencer F. Baird | 1957 | -43.72 | -107.6 | 10.99 | -3141 | Gravity core | 3-7 | 17 | 3 |
| 16370 | RV Spencer F. Baird | 1958 | -11.7 | -109.72 | 25.27 | -3296 | Gravity core | 0-3 | 1 | 1 |
| 16650 | RV Spencer F. Baird | 1958 | -18.33 | -79.34 | 20.07 | -3157 | Gravity core | 8-12 | 49 | 7 |
| 16651 | RV Spencer F. Baird | 1958 | -27.93 | -106.92 | 21.45 | -3039 | Gravity core | 8-15 | 67 | 6 |
| 16652 | RV Spencer F. Baird | 1958 | -27.15 | -109.83 | 21.45 | -2819 | Gravity core | 10-14 | 51 | 5 |
| 16653 | RV Spencer F. Baird | 1958 | -14.73 | -112.1 | 24.9 | -3034 | Gravity core | 12-15 | 69 | 7 |
| 16654 | RV Spencer F. Baird | 1958 | -9.88 | -110.68 | 25.26 | -2712 | Gravity core | 5-9 | 46 | 7 |
| 17162 | RV Vema | 1959 | 28.4 | -77.93 | 24.89 | -1004 | Piston core | 0-2 | 136 | 8 |
| 17359 | RV Vema | 1959 | -9.75 | -34.4 | 27.14 | -4123 | Piston core | 5-6 | 86 | 6 |

**Table S2:** Dissolution analysis: linear mixed-effects model (LMER, ANOVA), using size variation (95^th^ percentile of the population) as the response variable, species as the random effects and either a null model (H0) or water depth (H1) as the explanatory variable. Columns: model explanatory variables (fixed effects), degrees of freedom, Akaike Information Criterion, log-likelihood, model deviance, chi-squared, P value, marginal R squared.

| Exp. Var. | Df | AIC | logLik | Dev. | Chi-sq | P-value | R<sub>m</sub><sup>2</sup> |
| --- | --- | --- | --- | --- | --- | --- | --- |
| null | 5 | 151.67 | -70.84 | 141.67 | NA | NA | 0 |
| water depth | 6 | 151.75 | -69.87 | 139.74 | 1.93 | 0.16 | 0.00 |

### Museum bias analysis methods

to assess the size bias in the Buckley Collection, we re-sampled ten bulk sediments of the NHMUK Ocean-Bottom Deposits Collection (OBD) Collection, from which the Buckley Collection was created (Fig. 1a, Table S3). Samples were chosen to encompass different oceans, latitudes and marine expeditions; however, the final choice also depended on the availability of bulk sediment samples in the OBD Collection. Half of the amount available in the OBD containers was further split into two equal parts, leaving an archive sample and a sample to be processed. The sample processing consisted of weighing, wet washing over a 63μm sieve and drying in a 60°C oven. The residues were further dry sieved over a 150μm sieve and the coarser fraction was split with a microsplitter as many times as needed to produce a representative aliquot containing around 300 planktonic Foraminifera specimens (see Al-Sabouni et al. 2007). All specimens in each of the nine final splits were identified by MCR and MK under a stereomicroscope to species level, resulting in a total of 2,611 individuals belonging to 31 species (see also Rillo et al. 2019). We mounted species-specific slides from the resampled samples, calculated the relative abundance of each species in each sample, and then extracted shell size data in the same way as for the slides of the Buckley Collection. Only the sizes of species also present in the Buckley Collection samples were measured, resulting in 1824 specimens’ shell sizes from 20 species (Table S3). See Methods section in the main text for further information.

**Table S3:** Bias analysis: samples re-sampled from the Ocean-Bottom Deposits (OBD) Collection at the Natural History Museum, London (NHMUK) used in the museum collection size bias analysis. Columns: NHMUK Internal Record Number of the sediment in the OBD Collection; name of the Vessel that collected the sample; latitude (Lat) and longitude (Long) given in decimal degrees; water depth in meters; sampled mass in grams; number of times sediment was split N(splits) to achieve around 300 specimens; number of planktonic Foraminifera specimens N(ind) and species N(ssp) measured in each re-sampled sample.

| OBD_IRN | Vessel | Lat | Long | Depth | Mass (g) | N(splits) | N(ind) | N(ssp) |
| --- | --- | --- | --- | --- | --- | --- | --- | --- |
| 32657 | HMS *Challenger* | -50.01667 | 123.06667 | -3976 | 0.187 | 5 | 318 | 7 |
| 34991 | HMS *Challenger* | -21.25 | -14.03333 | -3740 | 9.35 | 7 | 265 | 21 |
| 33668 | HMS *Challenger* | -0.7 | 147 | -2213 | 1.98 | 7 | 331 | 16 |
| 33286 | HMS *Challenger* | 24.33333 | -24.46667 | -5153 | 2.73 | 5 | 260 | 18 |
| 34671 | HMS *Egeria* | -19.56667 | 64.63333 | -2708 | 1.23 | 5 | 376 | 24 |
| 34993 | HMS *Egeria* | -15.65 | -179.0625 | -2519 | 2.42 | 8 | 300 | 13 |
| 36053 | HMS *Penguin* | -26.93667 | 111.18167 | -3350 | 1.49 | 5 | 279 | 18 |
| 37148 | HMS *Sealark* | -7.59167 | 61.48333 | -3507 | 2.86 | 8 | 305 | 18 |
| 38482 | HMS *Waterwitch* | -40.45 | 49.81667 | -3780 | 1.51 | 6 | 177 | 11 |
| 14609 | Alpha 6 | 85.25 | -167.9 | -1774 | 0.57 | 4 | 226 | 1 |
|  |  |  |  |  |  | **TOTAL** | **2837** | **31** |

### Linear mixed-effects regression using the bias analysis data

using the re-sampled populations described above, we tested whether relative abundance variation significantly explains population shell size variation. Since the re-sampled data includes only ten samples (Fig. 1a), there is not enough data to run species-specific generalised linear models (GLM). Instead, we used linear mixed-effect models (LMER). The log-transformed 95^th^ percentiles of the population shell size distributions were modelled as the response variable, and the independent fixed effect was the species’ relative abundance in the re-sampled sample. Species were modelled as random effects, allowing for random intercepts and slopes (i.e., the intercept and slope of the relationship between shell size and the relative abundance may vary among species). We used the Likelihood Ratio Test (LRT) to compare the likelihood of the fixed effect. We calculated the LRT between the models with and without the effect. Significance of each fixed effect was given through the LRT. Marginal R^2^ (R_m_^2^), which is associated with the fixed effects, was calculated for each LMER model.

**Table S4:** Bias analysis: linear mixed-effects model (LMER, ANOVA) using the re-sampled data, and size variation (log-transformed 95^th^ percentile of the population) as the response variable, species as the random effects (r.e.) and either a null model (H0) or relative abundance (H1) as the explanatory variable. Columns: model explanatory variables (fixed effects), degrees of freedom, Akaike Information Criterion, log-likelihood, model deviance, chi-squared, P value, marginal R squared.

| Exp. Var. | Df | AIC | logLik | Dev. | Chi-sq | P-value | $R_m^2$ |
| --- | --- | --- | --- | --- | --- | --- | --- |
| null | 5 | 130.20 | -60.01 | 120.20 | NA | NA | 0 |
| relative abundance | 6 | 130.02 | -59.01 | 118.02 | 2.18 | 0.14 | 0.07 |

**Table S5:** Analysis considering relative abundance samples within a 300 km distance: model selection of the linear and quadratic models testing if planktonic foraminifera shell size (represented by the 95^th^ percentile of each population size distribution) can be predicted by mean annual sea-surface temperature (sst linear effect, sst^2^ quadratic effect), mean annual net-primary productivity (npp) and/or species’ relative abundance (abund, calculated as the median relative abundance of all samples within a 300 km distance of each morphometric sample). A model including the interaction between sst and npp (sst: npp) was also considered. Columns: species, explanatory variables, degrees of freedom, log-likelihood, Akaike Information Criterion corrected for small sample size (AICc), AICc difference between models (ΔAICc), model weight, adjusted R squared (values above 0.15 are in bold). All models within two ΔAICc units are shown and considered equally plausible.

| Species | Exp. Var. | df | logLik | AICc | ΔAICc | weight | R<sub>adj</sub><sup>2</sup> |
| --- | --- | --- | --- | --- | --- | --- | --- |
| *T. sacculifer* | sst | 3 | -7.04 | 20.79 | 0 | 0.38 | 0.25 |
| *T. sacculifer* | sst2 | 4 | -6.78 | 22.78 | 1.99 | 0.14 | 0.24 |
| *G. ruber* | null | 2 | -12.45 | 29.22 | 0 | 0.18 | 0 |
| *G. ruber* | sst + pp | 4 | -10.34 | 29.85 | 0.63 | 0.13 | 0.05 |
| *G. ruber* | sst | 3 | -11.61 | 29.9 | 0.67 | 0.13 | 0.02 |
| *G. ruber* | abund | 3 | -11.69 | 30.06 | 0.84 | 0.12 | 0.01 |
| *G. ruber* | pp | 3 | -12.01 | 30.7 | 1.47 | 0.09 | 0 |
| *G. ruber* | sst : pp | 5 | -9.45 | 30.71 | 1.49 | 0.08 | 0.07 |
| *G. globobatus* | sst : pp | 5 | 5.29 | 1.3 | 0 | 0.27 | 0.19 |
| *G. globobatus* | sst + pp | 4 | 3.4 | 2.4 | 1.1 | 0.16 | 0.13 |
| *G. globobatus* | sst2 : pp | 6 | 5.82 | 3.07 | 1.77 | 0.11 | 0.19 |
| *G. siphonifera* | sst + pp | 4 | -2.22 | 13.69 | 0 | 0.29 | 0.33 |
| *G. siphonifera* | sst | 3 | -4.24 | 15.2 | 1.51 | 0.14 | 0.27 |
| *G. siphonifera* | sst : pp | 5 | -1.63 | 15.2 | 1.52 | 0.14 | 0.33 |
| *N. dutertrei* | null | 2 | 3.59 | -2.73 | 0 | 0.22 | 0 |
| *N. dutertrei* | pp | 3 | 4.34 | -1.76 | 0.98 | 0.13 | 0.01 |
| *N. dutertrei* | abund | 3 | 4.17 | -1.42 | 1.31 | 0.11 | 0 |
| *N. dutertrei* | sst | 3 | 4.12 | -1.32 | 1.41 | 0.11 | 0 |
| *N. dutertrei* | sst2 | 4 | 5.25 | -0.9 | 1.83 | 0.09 | 0.04 |
| *P. obliquiloculata* | sst | 3 | 1.89 | 3.08 | 0 | 0.35 | 0.21 |
| *P. obliquiloculata* | sst + pp | 4 | 2.26 | 4.97 | 1.89 | 0.13 | 0.21 |
| *G. menardii* | sst | 3 | -2.53 | 12.02 | 0 | 0.18 | 0.06 |
| *G. menardii* | sst + abund | 4 | -1.29 | 12.26 | 0.23 | 0.16 | 0.1 |
| *G. menardii* | null | 2 | -3.92 | 12.29 | 0.27 | 0.16 | 0 |
| *G. menardii* | abund | 3 | -3.02 | 13.01 | 0.99 | 0.11 | 0.02 |
| *G. menardii* | pp | 3 | -3.48 | 13.92 | 1.9 | 0.07 | -0.01 |
| *G. truncatulinoidea* | sst2 + pp | 5 | 2.99 | 6.51 | 0 | 0.36 | 0.32 |
| *G. truncatulinoidea* | sst2 : pp | 6 | 3.88 | 7.9 | 1.39 | 0.18 | 0.34 |
| *G. inflata* | pp | 3 | 11.11 | -14.63 | 0 | 0.38 | 0.15 |
| *G. inflata* | null | 2 | 9.08 | -13.41 | 1.22 | 0.21 | 0 |

**Table S6:**
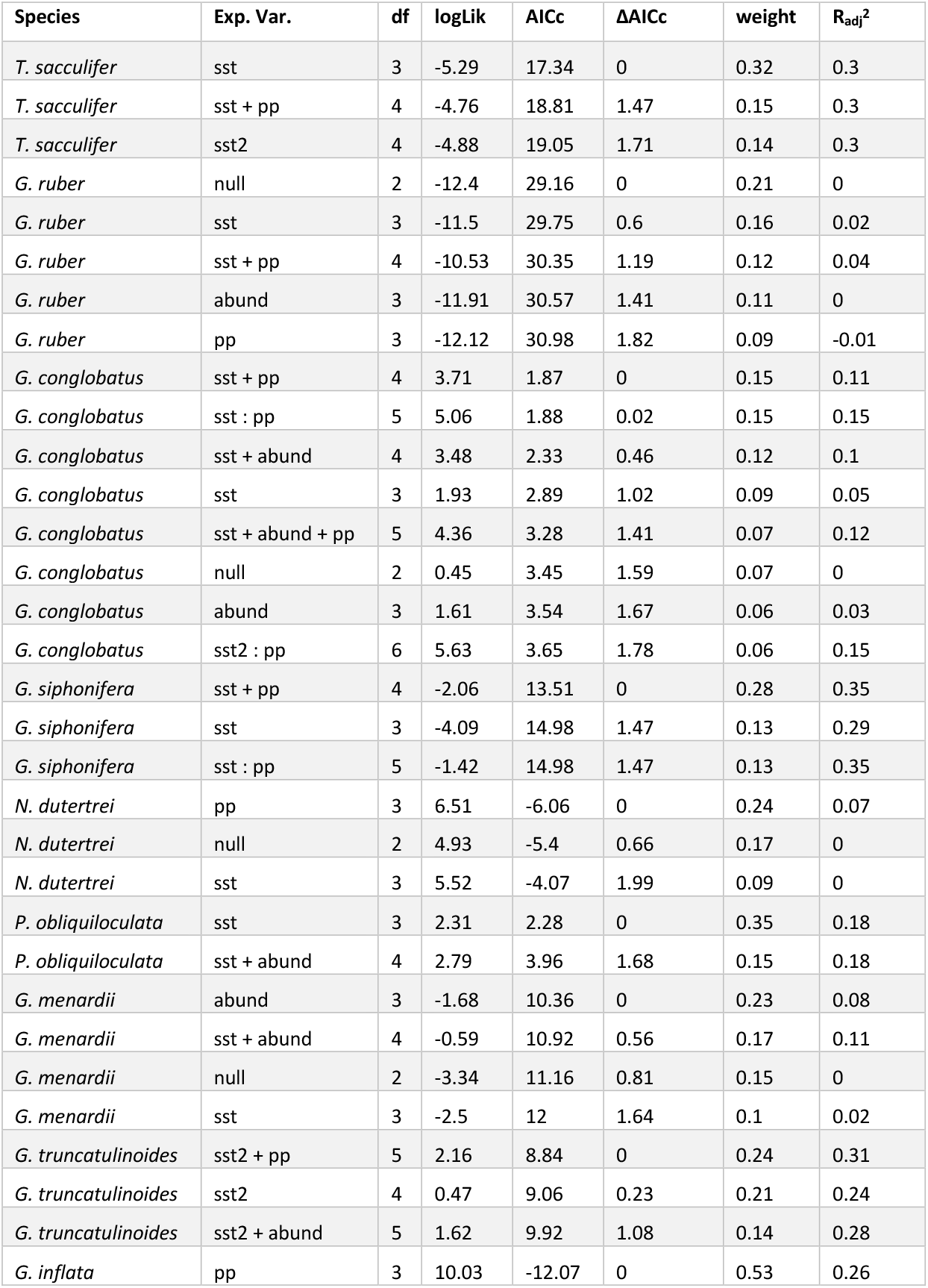
Analysis excluding historical samples collected by dredging the ocean floor, namely samples collected by HMNZS *Pukaki,* HMS *Challenger* (dredge only), HMS *Enterprise* and RNZFA *Tui* (see Table S1). Columns and variables: see description of Table S5.

**Fig. S1:**
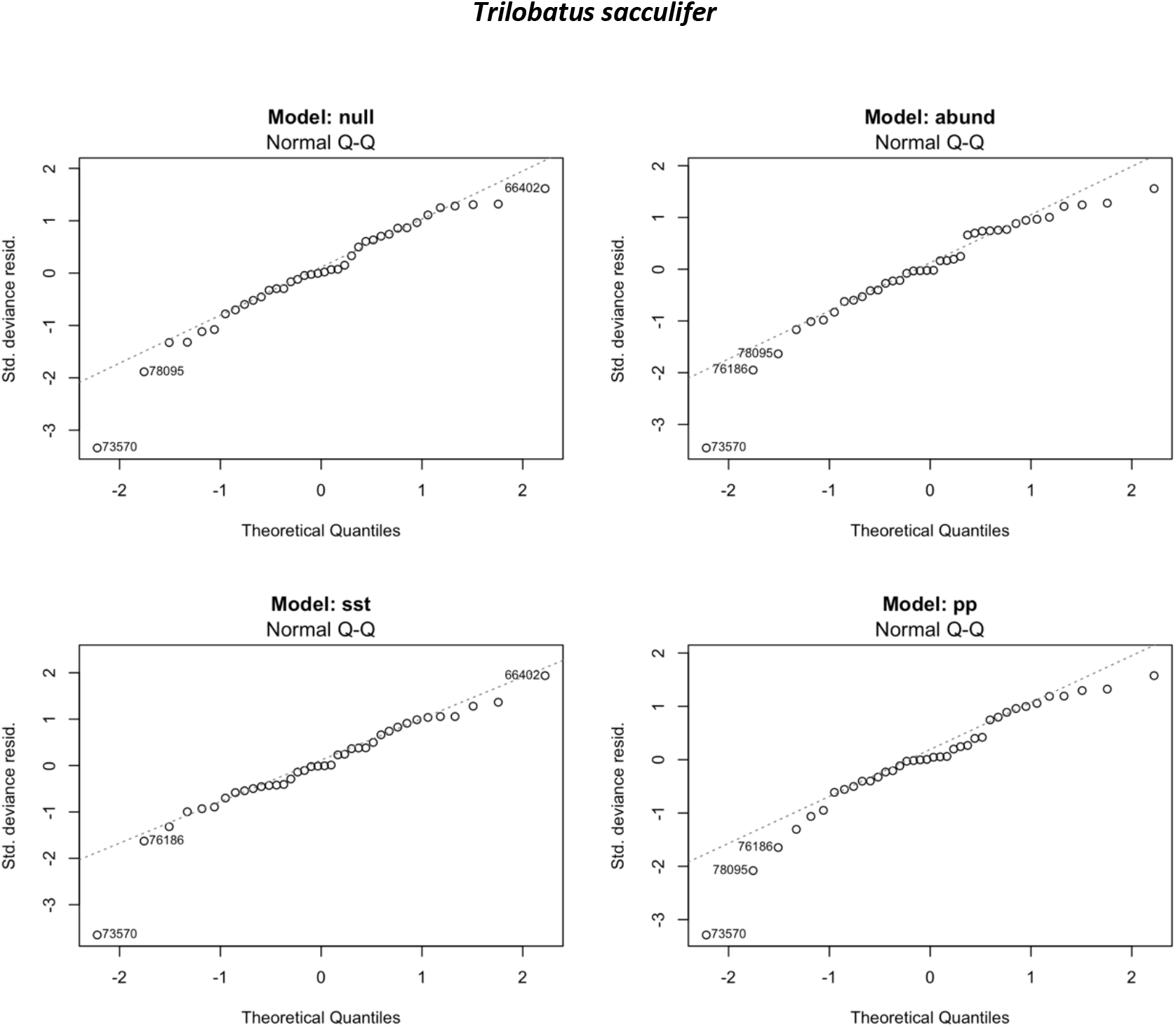
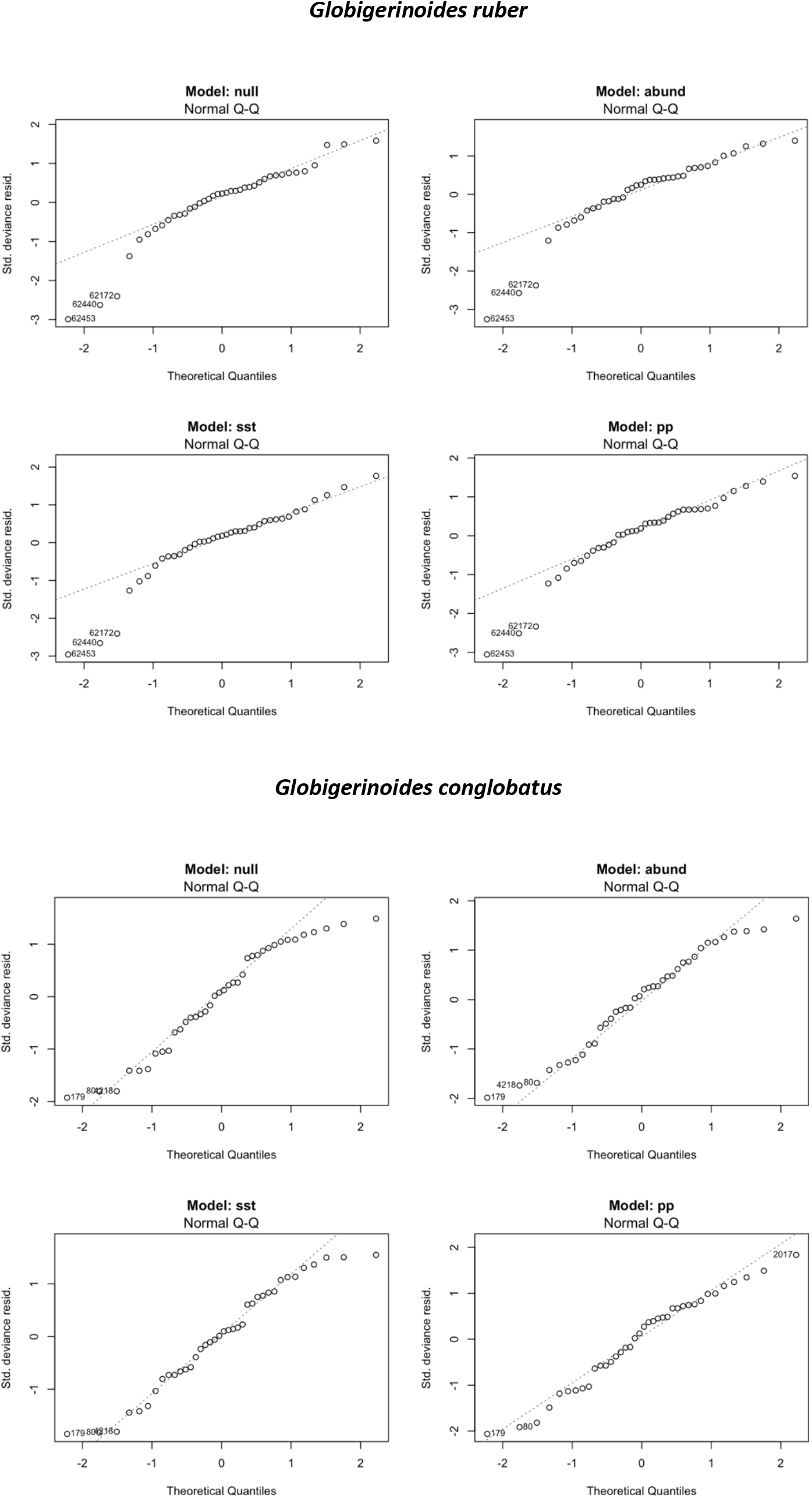
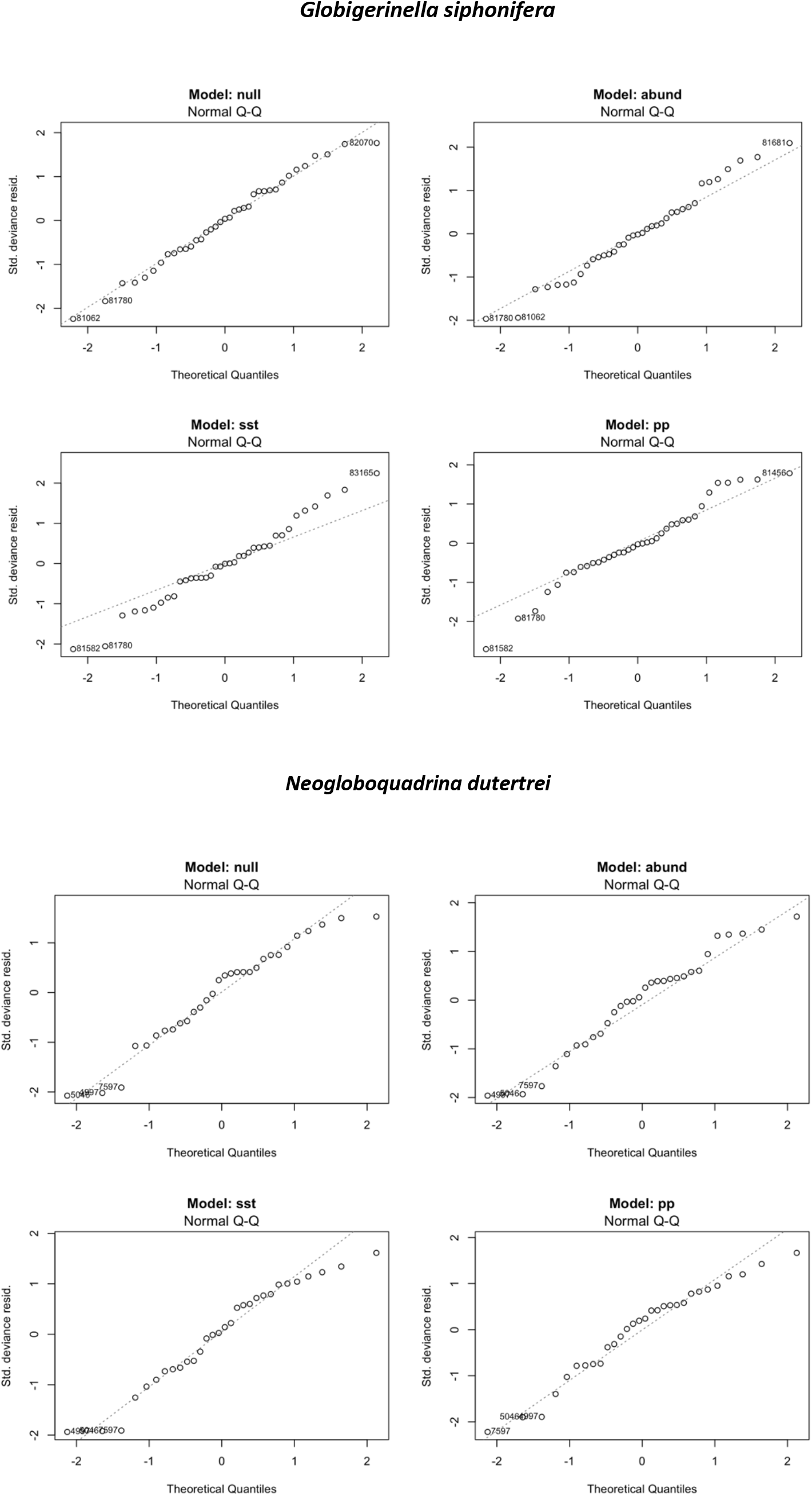
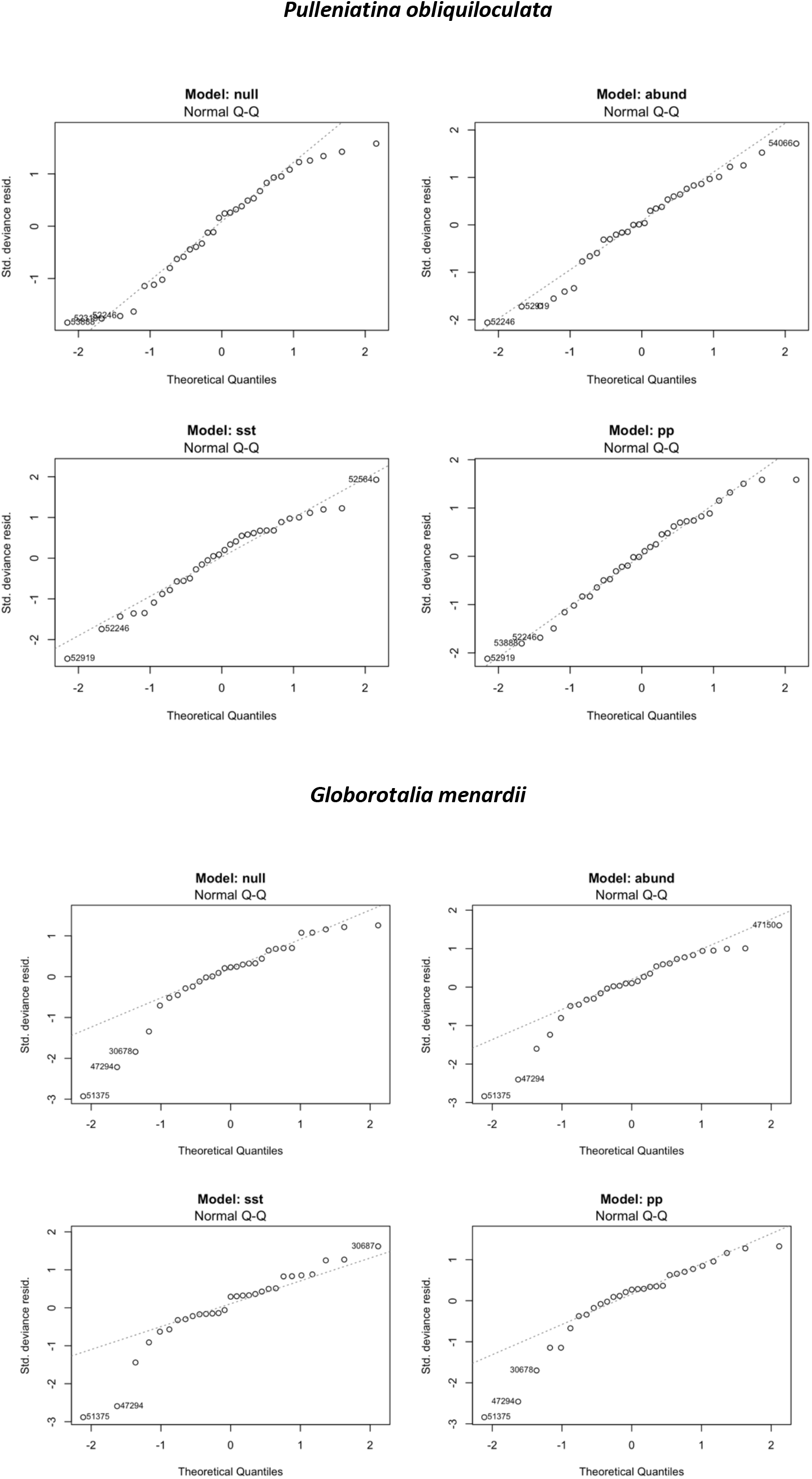
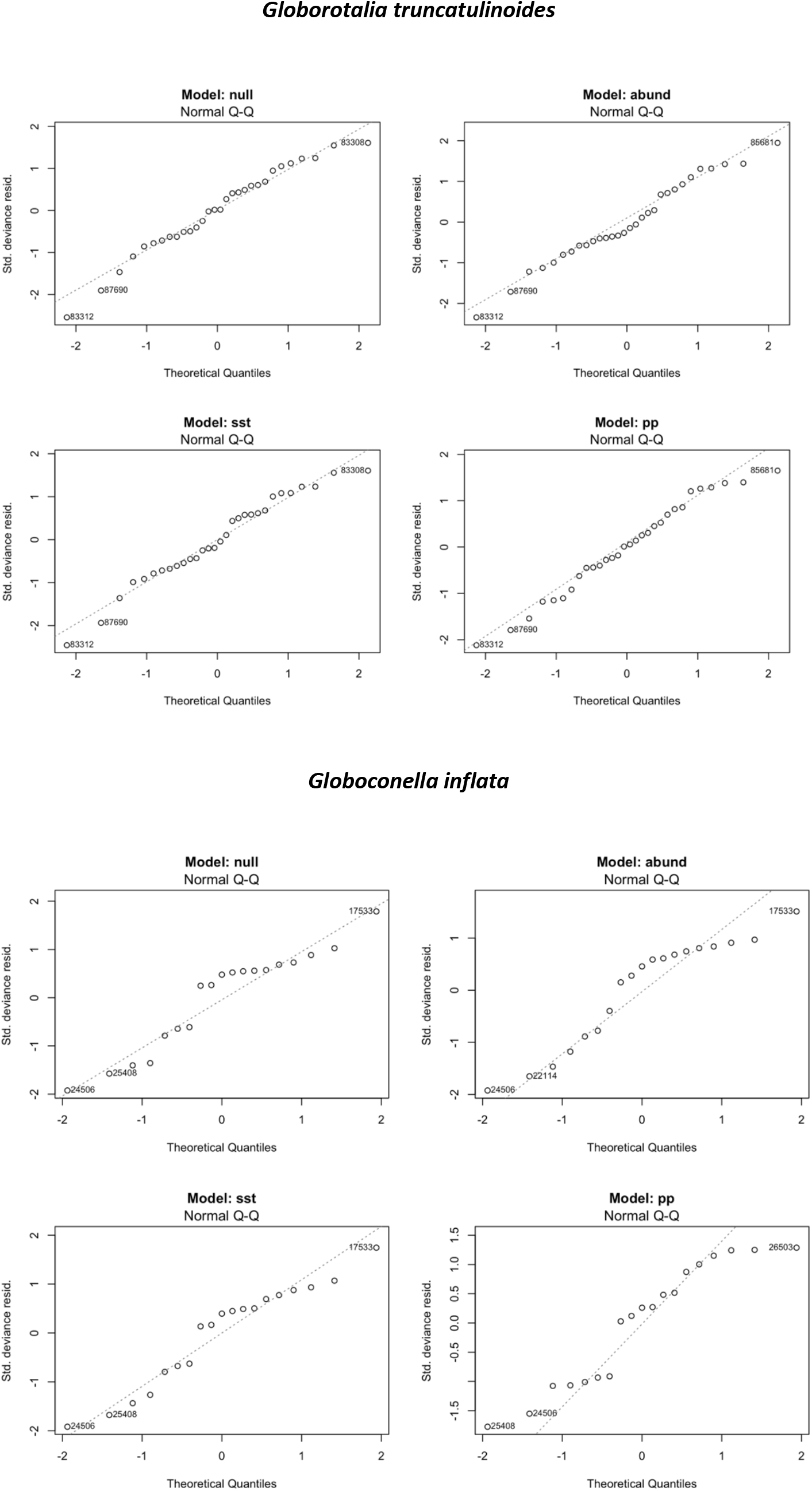
Residual plots of linear models per species. Models: null, abund (relative abundances), sst (mean annual sea surface temperature), and pp (mean annual net primary productivity).

